# Interhomolog polymorphism shapes meiotic crossover within *RAC1* and *RPP13* disease resistance genes

**DOI:** 10.1101/290478

**Authors:** Heïdi Serra, Kyuha Choi, Xiaohui Zhao, Alexander R. Blackwell, Ian R. Henderson

## Abstract

During meiosis chromosomes undergo DNA double-strand breaks (DSBs), which can produce crossovers via interhomolog repair. Meiotic recombination frequency is variable along chromosomes and concentrates in narrow hotspots. We mapped crossovers within *Arabidopsis thaliana* hotspots located within the *RAC1* and *RPP13* disease resistance genes, using varying haplotypic combinations. We observed a negative non-linear relationship between interhomolog divergence and crossover frequency, consistent with polymorphism suppressing crossover repair of DSBs. Anti-recombinase mutants *fancm*, *recq4a recq4b*, *figl1* and *msh2*, or lines with increased *HEI10* dosage, are known to show increased crossovers. Surprisingly, *RAC1* crossovers were either unchanged or decreased in these genetic backgrounds. We employed deep-sequencing of crossovers to examine recombination topology within *RAC1*, in wild type, *fancm* and *recq4a recq4b* mutant backgrounds. The *RAC1* recombination landscape was broadly conserved in anti-recombinase mutants and showed a negative relationship with interhomolog divergence. However, crossovers at the *RAC1* 5’-end were relatively suppressed in *recq4a recq4b* backgrounds, indicating that local context influences recombination outcomes. Our results demonstrate the importance of interhomolog divergence in shaping recombination within plant disease resistance genes and crossover hotspots.

## INTRODUCTION

Meiosis is a specialized cell division that is central to sexual reproduction in eukaryotes (Villeneuve & Hillers, 2001; Mercier *et al*, 2015). It is characterized by a single round of DNA replication, followed by two successive rounds of chromosome segregation, generating four haploid gametes from a single diploid mother cell (Villeneuve & Hillers, 2001; Mercier *et al*, 2015). During prophase I, homologous chromosomes pair and undergo reciprocal genetic exchange, termed crossover (Hunter, 2015). Crossovers ensure accurate chromosome segregation, by creating a physical link between homologous chromosomes that promotes balanced segregation during the first meiotic division (Villeneuve & Hillers, 2001; Mercier *et al*, 2015). Importantly, meiotic crossovers also create genetic diversity by recombining linked variation (Villeneuve & Hillers, 2001; Barton & Charlesworth, 1998; Mercier *et al*, 2015). Meiotic recombination thus impacts upon genetic adaptation in sexual populations, by combining independently arising mutations more rapidly than in asexual species (Barton and Charlesworth, 1998).

Meiotic recombination initiates via DNA double-strand breaks (DSBs) generated by SPO11 topoisomerase VI-related transesterases (Szostak *et al*, 1983; Keeney *et al*, 1997; de Massy, 2013). In Arabidopsis ∼100-200 meiotic DSBs form per meiosis, estimated from immunostained RAD51, DMC1, RPA1 and _Ɣ_H2A.X foci that occur along paired chromosomes at leptotene stage (Ferdous *et al*, 2012; Yelina *et al*, 2015; Chelysheva *et al*, 2010). In budding yeast, endonuclease and exonuclease activities (Mre11-Rad50-Xrs2, Sae2 and Exo1) act at DSB sites to generate 3’-overhanging single-strand DNA (ssDNA) (Neale *et al*, 2005; Cannavo & Cejka, 2014; Zakharyevich *et al*, 2010; Garcia *et al*, 2011), between ∼100s and 1000s of nucleotides in length (Mimitou *et al*, 2017; Sun *et al*, 1991). Resected ssDNA is bound first by RPA1 and then RAD51 and DMC1 proteins, which together promote interhomolog invasion and formation of a displacement loop (D-loop) (Brown & Bishop, 2015). Stabilization of the D-loop involves template-driven DNA synthesis from the invading 3’-end (Keeney & Neale, 2006; Hunter, 2015). Strand invasion intermediates can then undergo second-end capture to form double Holliday junctions (dHJ), which can be resolved as a crossover or non-crossover, or dissolved (Kohl & Sekelsky, 2013; Schwacha & Kleckner, 1994).

The conserved ZMM pathway acts to promote meiotic DSB repair via dHJs and crossover formation (Lynn *et al*, 2007; Hunter, 2015; Mercier *et al*, 2015). In Arabidopsis ∼10 DSBs per meiosis are repaired as crossovers (Wijnker *et al*, 2013; Copenhaver *et al*, 1998; Salomé *et al*, 2012; Giraut *et al*, 2011). The majority (∼90%) of these crossovers are dependent on the ZMM pathway in Arabidopsis (Mercier *et al*, 2015). This pathway includes ZIP4/SPO22, the SHOC1 XPF endonuclease and its interacting partner PTD, the MER3 DNA helicase, the HEI10 E3 ligase, the MSH4/MSH5 MutS-related heterodimer and the MLH1/MLH3 MutL-related heterodimer (Lynn *et al*, 2007; Mercier *et al*, 2015). ZMM factors are thought to stabilise interhomolog joint molecules, including dHJs, and promote crossover resolution (De Muyt *et al*, 2018). ZMM-dependent crossovers (also known as class I) show the phenomenon of interference, meaning that they are more widely distributed than expected at random (Lynn *et al*, 2007; Mercier *et al*, 2015; Kleckner *et al*, 2004; Berchowitz & Copenhaver, 2010).

In plants and other eukaryotes the large excess of initiating meiotic DSBs proceed to resection and strand invasion, but are repaired as non-crossovers (detectable as gene conversions), or via inter-sister repair (Mercier *et al*, 2015). Dissolution of strand invasion events occurs via partially redundant anti-crossover pathways in Arabidopsis that include, (i) the FANCM helicase and its cofactors MHF1 and MHF2 (Crismani *et al*, 2012; Girard *et al*, 2014; Knoll *et al*, 2012), (ii) the BTR complex: RECQ4A, RECQ4B, TOPOISOMERASE3a and RECQ4-MEDIATED INSTABILITY1 (RMI1) (Séguéla-Arnaud *et al*, 2016, 2015; Bonnet *et al*, 2013; Hartung *et al*, 2007; Higgins *et al*, 2011), and (iii) FIDGETIN-LIKE1 (FIGL1) and FLIP1 (Girard *et al*, 2015; Fernandes *et al*, 2017a). Plants mutated in these anti-crossover pathways show increased non-interfering crossovers, which are also known as class II events (Mercier *et al*, 2015). This likely occurs as a consequence of reduced dissolution of interhomolog strand invasion events, which are alternatively repaired by the non-interfering crossover pathway(s) (Crismani *et al*, 2012; Girard *et al*, 2015; Séguéla-Arnaud *et al*, 2015), including MUS81 (Berchowitz *et al*, 2007; Higgins *et al*, 2008). Hence, alternative repair pathways act on SPO11-dependent DSBs during meiosis to balance crossover and non-crossover outcomes.

Due to the formation of interhomolog joint molecules during recombination, sequence polymorphisms between recombining chromosomes can result in mismatched base pairs (Chakraborty & Alani, 2016). During the mitotic cell cycle DNA mismatches or short insertion-deletions (indels) caused by base mis-incorporation during replication, or exogenous DNA damage, can be detected by MutS-related heterodimers (Harfe & Jinks-Robertson, 2000). MutS recognition of mismatches and the subsequent promotion of repair plays a major anti-mutagenic role *in vivo* (Harfe & Jinks-Robertson, 2000). MutS complexes also play anti-crossover roles during meiosis when heterozygosity leads to sequence mismatches, following interhomolog strand invasion (Alani *et al*, 1994; Hunter *et al*, 1996; Emmanuel *et al*, 2006). Accumulating evidence also indicates that class I and II crossover repair pathways show differential sensitivity to levels of interhomolog polymorphism. For example, Arabidopsis *fancm* mutations show increased crossovers in inbred, but not in hybrid contexts, whereas *figl1* and *recq4a recq4b* mutations are effective at increasing crossovers in both inbreds and in hybrids (Ziolkowski *et al*, 2015; Girard *et al*, 2015; Fernandes *et al*, 2017b; Séguéla-Arnaud *et al*, 2015; Serra *et al*, 2018; Ziolkowski *et al*, 2017). This implies that the non-interfering crossover repair pathways acting in these backgrounds are influenced differently by interhomolog polymorphism. Genome-wide mapping of crossovers in anti-crossover mutants, or backgrounds with additional copies of the ZMM gene *HEI10*, have also shown that the resulting recombination increases are highly distalized towards the sub-telomeres, correlating with regions of lowest interhomolog polymorphism (Ziolkowski *et al*, 2017; Serra *et al*, 2018; Fernandes *et al*, 2017b). This may reflect a preference for recombination pathways to act in less divergent regions. At larger physical scales structural rearrangements, such as translocations and inversions, are potently associated with crossover suppression (Fransz *et al*, 2016; Fang *et al*, 2012).

In this work we explore the influence of interhomolog polymorphism on meiotic recombination at the scale of crossover hotspots in *Arabidopsis thaliana*. Specifically, we map crossovers across the *RAC1* and *RPP13* disease resistance genes, which encode proteins that recognise effector proteins from the oomycete pathogens *Albugo candida* and *Hylaoperonospora parasitica*, respectively (Borhan *et al*, 2004; Allen *et al*, 2004). We observe a strong non-linear negative relationship between interhomolog polymorphism and crossover frequency at both *RAC1* and *RPP13*, supporting a local inhibitory effect of heterozygosity on crossover maturation. This relationship is observed using different *RAC1* haplotypic combinations, which vary in polymorphism density and pattern. Despite recombination rates increasing genome-wide in anti-crossover mutants and *HEI10* transgenics, *RAC1* crossover frequency was stable or significantly decreased in these backgrounds. Using deep sequencing of *RAC1* crossover molecules we show that the negative relationship between crossovers and interhomolog divergence is maintained in *fancm recq4a recq4b* anti-crossover mutants. However, crossover frequency in the 5’ region of *RAC1* was decreased in *recq4a recq4b* mutant backgrounds. The resistance of *RAC1* to genome-wide crossover increases may relate to the gene’s high level of interhomolog polymorphism, pericentromeric location and local chromatin environment.

## RESULTS

### Meiotic recombination and chromatin at *RAC1* and *RPP13* disease resistance genes

We previously identified the *RESISTANCE TO ALBUGO CANDIDA1* (*RAC1*) Arabidopsis disease resistance gene region as containing crossover hotspots, using both historical linkage disequilibrium (LD) estimates and experimental pollen-typing in Col×Ler F_1_ hybrids (Choi *et al*, 2013, 2016). *RAC1* encodes a TIR-NBS-LRR domain resistance protein, which recognises effectors from the oomycete pathogens *Albugo candida* and *Hylaoperonospora parasitica* (Borhan *et al*, 2004, 2001; Speulman *et al*, 1998). *RAC1* exists as a singleton TIR-NBS-LRR gene in most accessions and shows high levels of population genetic diversity (e.g. θ=0.012/0.013; π=0.043/0.054) (Choi *et al*, 2013; Long *et al*, 2013; Cao *et al*, 2011; Choi *et al*, 2016). We compared the *RAC1* locus to a recombination map of 3,320 crossovers mapped by genotyping-by-sequencing (GBS) of 437 Col×Ler F_2_ individuals (mean resolution of crossovers=970 bp) (Fig. 1A) (Serra *et al*, 2018; Choi *et al*, 2016). We also assessed levels of interhomolog polymorphism by measuring the density of Col/Ler SNPs per 100 kb (Zapata *et al*, 2016), in addition to levels of DNA methylation as an indication of heterochromatin (Fig. 1A) (Stroud *et al*, 2013). Together this showed that *RAC1* is located on the edge of pericentromeric heterochromatin, in a region of higher than average crossover frequency and interhomolog polymorphism (Fig. 1A).

**Figure 1.**
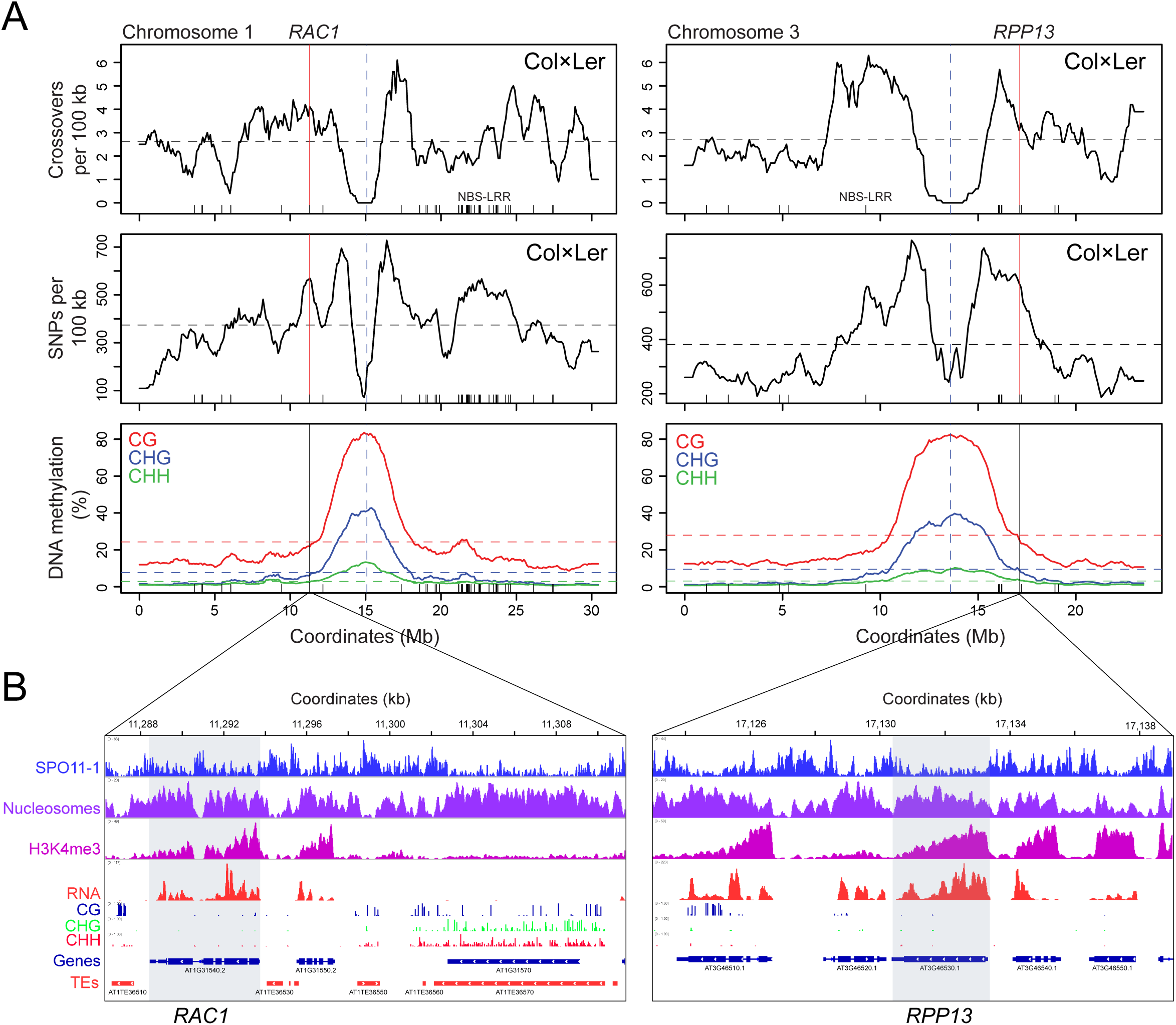
Chromatin and recombination landscapes around the *RAC1* and *RPP13* NBS-LRR disease resistance genes. **(A)** Crossover frequency (crossovers/100 kb mapped by genotyping-by-sequencing of Col×Ler F_2_) (Choi *et al*, 2016; Serra *et al*, 2018), interhomolog divergence (Col/Ler SNPs/100 kb) (Zapata *et al*, 2016), and % DNA methylation (CG red, CHG blue, CHH green) (Stroud *et al*, 2013), along chromosomes 1 and 3. Vertical dotted lines indicates the centromeres. Mean values are indicated by horizontal dotted lines. NBS-LRR gene positions are indicated by ticks on the x-axis. The position of *RAC1* and *RPP13* are indicated by the solid vertical lines. **(B)** Histograms for the *RAC1* and *RPP13* regions showing library size normalized coverage values for SPO11-1-oligonucleotides (blue), nucleosome occupancy (purple, MNase-seq), H3K4me3 (pink, ChIP-seq), RNAseq (red) and % DNA methylation (BSseq) in CG (blue), CHG (green) and CHH (red) sequence contexts (Choi *et al*, 2018; Stroud *et al*, 2013). Gene (blue) and transposon (red) annotations are highlighted, and the positions of *RAC1* and *RPP13* are indicated by grey shading.

Using historical recombination maps generated by analysing the 1,001 Genomes Project SNP data, we identified *RPP13* as a further NBS-LRR gene with higher than average historical crossover frequency (10.56/10.57 cM/Mb), and high levels of population SNP diversity (θ=0.011/0.013, π=0.044/0.045) (Choi *et al*, 2013; Long *et al*, 2013; Cao *et al*, 2011; Choi *et al*, 2016). These levels of diversity and recombination were comparable to those observed at *RAC1*. RPP13 recognizes the *Hylaoperonospora parasitica* effector ATR13 to mediate disease resistance, and which together display co-evolutionary dynamics (Allen *et al*, 2004; Rose *et al*, 2004). Similar to *RAC1*, *RPP13* is a singleton NBS-LRR gene located on the edge of pericentromeric heterochromatin, in a region of higher than average crossover frequency and interhomolog polymorphism (Fig. 1A).

We next examined the *RAC1* and *RPP13* regions using genome-wide maps of chromatin and meiotic recombination (Choi *et al*, 2016, 2018; Stroud *et al*, 2013). Nucleosome occupancy was assessed using MNase-seq data and observed to be enriched within the gene exons and depleted within intergenic promoter, intron and terminator regions (Fig. 1B). SPO11-1-oligonucleotides mark meiotic DSB sites and show an inverse pattern to nucleosome occupancy (Choi *et al*, 2018). Consistently, at *RAC1* and *RPP13* we observed higher levels of SPO11-1-oligonucleotides in nucleosome-depleted intergenic regions (Fig. 1B). H3K4me3 ChIP-seq showed enrichment at the 5’-end of the genes, consistent with active transcription (Zhang *et al*, 2009). Indeed, we observed evidence of *RAC1* and *RPP13* transcription using RNA-seq data from stage 9 flowers (Fig. 1B) (Choi *et al*, 2016, 2018). Both *RAC1* and *RPP13* show low levels of DNA methylation, in contrast to the *ATENSPM3* EnSpm/CACTA (AT1TE36570) element adjacent to *RAC1*, which is densely cytosine methylated, nucleosome-dense and suppressed for SPO11-1-oligos (Fig. 1B). The *RAC1* promoter intergenic region contains short fragments of several transposable elements, including *HELITRONY3* and *ATREP15* Helitrons (Fig. 1B). Transposable elements in these families have low DNA methylation, are nucleosome-depleted and show higher levels of SPO11-1-oligos, compared with other repeat families (Fig. 1B) (Choi *et al*, 2018). Therefore, despite the location of *RAC1* and *RPP13* on the edge of pericentromeric heterochromatin, these genes display euchromatic chromatin and recombination features (Fig. 1A-1B) (Stroud *et al*, 2013; Choi *et al*, 2018).

### Crossover hotspots within the *RAC1* and *RPP13* disease resistance genes

In order to experimentally measure crossover frequency and patterns within *RAC1* and *RPP13* we used pollen-typing (Choi *et al*, 2017; Drouaud & Mézard, 2011). This method uses allele-specific PCR amplification from F_1_ hybrid genomic DNA extracted from gametes, in order to quantify and sequence crossover molecules (Fig. 2A and Fig. S1) (Choi *et al*, 2017; Drouaud & Mézard, 2011). This method is directly analogous to mammalian sperm-typing methods (Kauppi *et al*, 2009; Cole *et al*, 2010; Baudat & de Massy, 2007; Tiemann-Boege *et al*, 2006). Genomic DNA is extracted from pollen (male gametophytes) collected from individuals that are heterozygous over a known crossover hotspot (Fig. 2A). Allele-specific primers annealing to polymorphic sites flanking the region of interest are used to PCR amplify single crossover or parental molecules, using diluted DNA samples (Fig. 2A). Titration is used to estimate the concentrations of amplifiable crossover and parental molecules. These values are used to calculate genetic distance (cM = (crossovers/(crossovers+parentals))×100) (Fig. 2A). Sanger sequencing of PCR products amplified from single crossover molecules is performed to identify internal recombination points, to the resolution of individual polymorphisms (Fig. 2A). Together this information describes the recombination rate (cM/Mb) topology throughout the PCR amplified region (Drouaud & Mézard, 2011; Choi *et al*, 2017). It is also possible to mass amplify crossover molecules, which may be pooled and then used with high throughput paired-end sequencing to identify crossover locations (Fig. 2A) (Choi *et al*, 2017).

**Figure 2.**
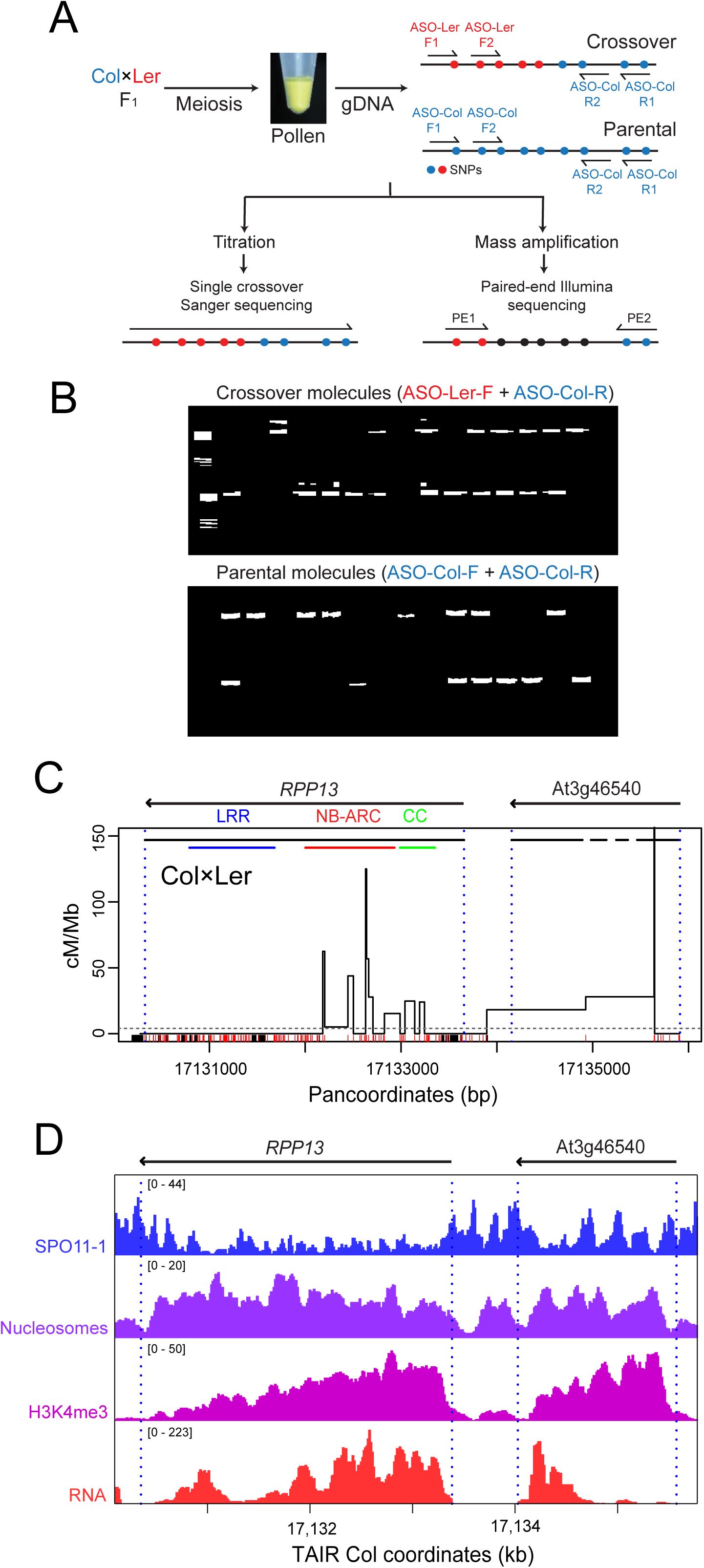
Crossover hotspots within the *RPP13* disease resistance gene. **(A)** Schematic of the pollen-typing method using allele-specific PCR to quantify and sequence crossover molecules. **(B)** Inset are representative ethidium bromide gels showing crossover and parental molecule *RAC1* PCR amplification products from diluted pollen F_1_ Col/Ler genomic DNA. **(C)** Crossover frequency (cM/Mb) within the region of the disease resistance gene *RPP13* measured using titration and sequencing of individual crossover molecules from Col×Ler pollen F_1_ genomic DNA. Gene TSS and TTS are indicated by vertical dotted blue lines. Horizontal lines indicate exon (black) positions, in addition to protein domains (coiled coil (green), NB-ARC (red) and LRR (blue)) for RPP13. Col/Ler SNPs (red) and indels (black) are indicated on the x-axis. Data is plotted against the Col/Ler panmolecule, which includes all insertions and deletions. The horizontal dotted line indicates the genome-average recombination rate for male Col×Ler crosses (Drouaud *et al*, 2007). **(D)** Histograms for the *RPP13* region showing library size normalized values for SPO11-1-oligonucleotides (blue), nucleosome occupancy (purple, MNase-seq), H3K4me3 (pink, ChIP-seq) and RNA-seq (red) (Choi *et al*, 2016, 2018).

To investigate whether *RPP13* was associated with crossover hotspots we designed and optimised Col/Ler allele-specific oligonucleotides flanking this resistance gene (Fig. S1). We performed DNA titrations to quantify crossover and parental molecules across *RPP13* and observed a genetic distance of 0.055 cM, equivalent to 9.78 cM/Mb across the 5,626 bp amplicon (Table S1). To analyse Sanger sequenced crossovers we plot their frequency against panmolecules, where we include all bases from both accessions (Fig. S2 and Tables S2-S5). For example, the *RPP13* amplicon is 5,431 bp in Col, 5,526 bp in Ler and 5,626 bp in the Col/Ler panmolecule, with 195 inserted bases from Ler and 100 from Col (Fig. S2 and Table S5). We sequenced 44 single crossover molecules and observed clustering of recombination events at the 5’-end of *RPP13*, overlapping regions encoding the coiled-coil and NB-ARC domains (Fig. 2C). *RPP13* shows a peak crossover rate of 125 cM/Mb (Table S6), compared to the genome average of 4.82 cM/Mb for male Col/Ler F_1_ hybrids (Drouaud *et al*, 2007). Crossovers were also observed in the adjacent gene At3g46540 (Fig. 2C). One crossover was observed in a single 5 bp interval within At3g46540, which results in a very high recombination estimate (Table S6). However, this likely reflects a sampling artefact, rather than the presence of a true hotspot. The region of highest crossover activity within *RPP13* overlaps with nucleosome-occupied, H3K4me3-modified exon sequences (Fig. 2D). In contrast, highest SPO11-1-oligos occur in flanking nucleosome-depleted intergenic regions (Fig. 2D).

The *RAC1* gene is located within a 9,482 bp (Col/Ler) pollen-typing PCR amplicon (Fig. 3). We previously reported analysis of 181 single crossover molecules within the *RAC1* amplicon (Choi *et al*, 2016), which we combined with an additional 59 events here to give a total of 240 crossovers (Table S8). We observed a peak rate of 61.7 cM/Mb within *RAC1* (Fig. 3A and Table S8). An adjacent gene contained within the amplicon, At1g31550 (*GDSL*), also showed intragenic crossovers (Fig. 3A) (Choi *et al*, 2016). Similar to *RPP13*, elevated crossover frequency within *RAC1* overlapped nucleosome-occupied and H3K4me3-enriched exon sequences (Fig. 3B) (Choi *et al*, 2018). Highest crossover frequency occurred within the *RAC1* 5′ exons encoding the NB-ARC and TIR domains (Fig. 3B). A further similarity with *RPP13*, is that highest levels of SPO11-1-oligos are observed in the nucleosome-depleted intergenic regions flanking *RAC1*, in addition to the largest intron (Figs. 2-3). Hence, both *RPP13* and *RAC1* have highest crossover frequency within transcribed gene 5′ regions, despite higher levels of initiating SPO11-1 DSBs occurring in the adjacent intergenic regions.

**Figure 3.**
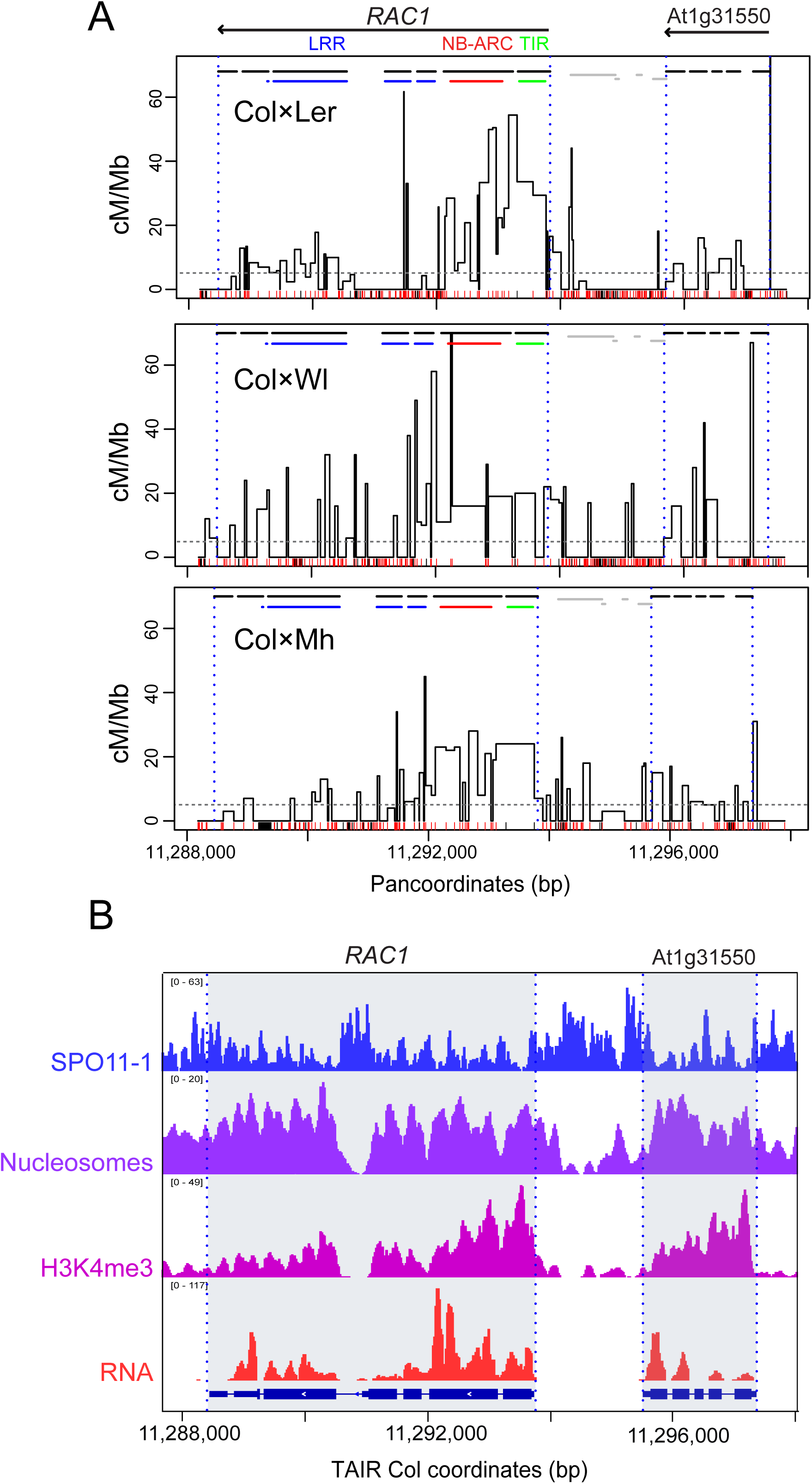
*RAC1* crossover hotspots in Col×Ler, Col×Wl and Col×Mh hybrids. **(A)** Crossover frequency (cM/Mb) within the region of the *RAC1* disease resistance gene measured using titration and sequencing of single crossover molecules from Col×Ler (0.074 cM), Col×Wl (0.074 cM) and Col×Mh (0.064 cM) pollen F_1_ genomic DNA. Recombination is plotted against the panmolecules, which include all bases from both parental accessions. Gene TSS/TTS are indicated by vertical dotted lines and exons by horizontal black lines. The position of RAC1 TIR (green), NB-ARC (red) and LRR (blue) domain-encoding sequences are indicated by the colored horizontal lines. SNPs (red) and indels (black) are indicated by the ticks on the x-axis. The horizontal dotted line indicates the genome-average recombination rate from male Col×Ler crosses (Drouaud *et al*, 2007). **(B)** Histograms for the *RAC1* region showing library size normalized values for SPO11-1-oligonucleotides (blue), nucleosome occupancy (purple, MNase-seq), H3K4me3 (pink, ChIP-seq) and RNA-seq (red) (Choi *et al*, 2018, 2016; Stroud *et al*, 2013). The positions of *RAC1* and *GDSL* (At1g31550) are indicated by grey shading.

### Interhomolog divergence suppresses crossovers within *RAC1* and *RPP13*

*RAC1* and *RPP13* show high levels of interhomolog polymorphism between Col and Ler, with 27.4 and 34.5 SNPs/kb, respectively (compared to the genome average of 3.85 SNPs/kb) (Zapata *et al*, 2016). This is also reflected in high levels of population genetic diversity at *RAC1* and *RPP13* (Choi *et al*, 2013; Long *et al*, 2013; Cao *et al*, 2011; Choi *et al*, 2016). We repeated *RAC1* pollen-typing with additional crosses using different parental accessions, where the pattern of interhomolog divergence varied, in order to investigate how patterns of interhomolog divergence influence local crossover frequency (Fig. 3A). Pollen-typing relies on allele-specific primers that anneal at SNPs or indels (Khademian *et al*, 2013; Choi *et al*, 2017). Therefore, we used the 1,001 Genomes Project data to identify accessions sharing the Col/Ler allele-specific primer polymorphisms, but differing with respect to internal polymorphisms within the *RAC1* amplicon (Fig. 3A and Fig. S2). This identified Mh-0 (Mühlen, Poland) and WI (Wildbad, Germany) as meeting these criteria. Col×Wl and Col×Mh have 33.0 and 21.1 SNPs/kb within the *RAC1* amplicon, respectively. We extracted pollen genomic DNA from Col×Wl and Col×Mh F_1_ hybrids and amplified and sequenced 92 and 124 crossover molecules, respectively (Fig. 3A). For Col×Ler and Col×Mh we performed DNA titration experiments and did not observe a significant difference in crossover frequency (*P=*0.309) (Table S7).

Crossover topology within the *RAC1* amplicon was conserved between the three haplotype combinations tested (Fig. 3A and Tables S8-S10). For instance, by comparing crossovers in adjacent 500 bp windows (counted against the Col reference sequence) we observed significant positive correlations between the recombination maps (Spearman’s Col×Ler vs Col×Wl *r*=0.595 *P=*9.14×10^-3^; Col×Ler vs Col×Mh *r*=0.612 *P=*6.91×10^-3^; Col×Wl vs Col×Mh *r*=0.723 *P=*6.96×10^-4^). For each cross, the highest crossover frequency was observed within the *RAC1* and *GDSL* transcribed regions (Fig. 3A and Tables S8-S10). In each case, we calculated the number of crossovers and polymorphisms in adjacent 500 bp windows (Fig. 4 and Table S11). SNPs were counted as one and indels were counted according to their length. In all cases, a significant negative relationship was observed between crossovers and polymorphisms (all *RAC1* windows, Spearman’s *r*=-0.685 *P*=1.11×10^-8^) (Fig. 4 and Table S11). This was also seen when analysing the *RPP13* Col×Ler data in the same manner (Spearman’s *r*=-0.890, *P*=2.43×10^-4^) (Fig. 4E and Table S12). We fitted a non-linear model to the data using the formula y = log(a)+b×x^(-c), where y is the number of crossovers, x is polymorphisms, a is the intercept and b and c are scale parameters. Together this shows a strong, negative, non-linear association between interhomolog polymorphism density and crossover frequency within *RAC1* and *RPP13*.

**Figure 4.**
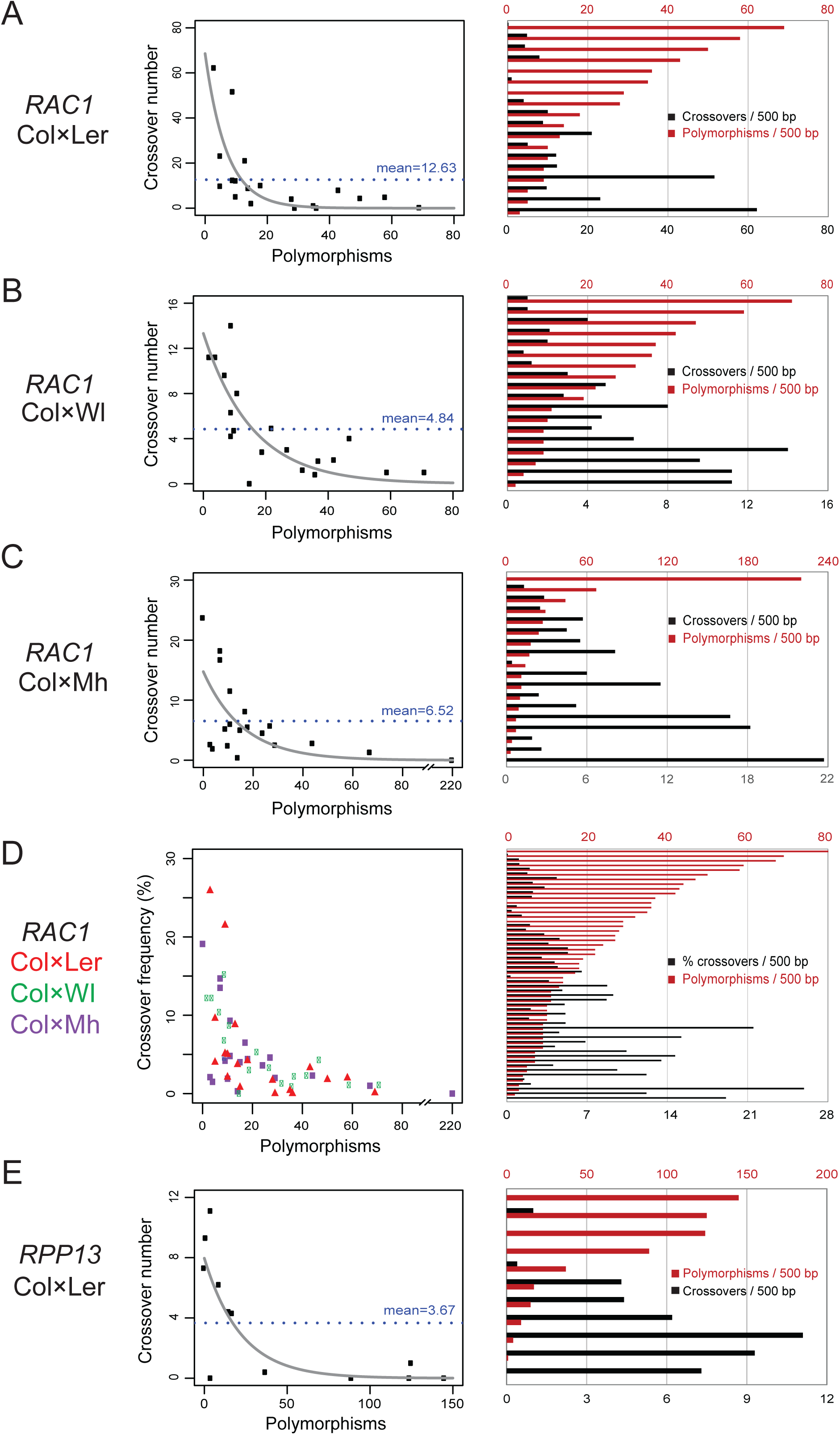
Interhomolog divergence suppresses crossovers within *RAC1* and *RPP13*. (A) Col×Ler crossovers and polymorphisms were calculated in adjacent 500 bp windows throughout the *RAC1* region, where SNPs are counted as one and indels by their length, using panmolecule coordinates. The blue horizontal dotted lines indicate the value of crossovers per window if they were evenly distributed. The grey line represents a non-linear model fitted to the data using the formula; y = log(a)+b×x^(-c), where y is the number of crossovers, x is polymorphisms, a is the intercept and b and c are scale parameters. On the right the same data are plotted as histograms of crossover (black) and polymorphisms (red) per 500 bp window. (B) As for (A), but analysing Col×Wl. (C) As for (A), but analysing Col×Mh. (D) As for (A), but analysing all windows from Col×Ler (red), Col×Wl (green) and Col×Mh (purple). Due to total crossovers analysed varying between hybrids, crossovers were first calculated as a % for each window. (E) As for (A), but analysing the *RPP13* amplicon from a Col×Ler hybrid.

Interestingly, the *RAC1* promoter region, which contains transposon fragments, shows higher recombination in Col×Mh compared with Col×Ler (3.41 versus 0.61 cM/Mb, respectively). This correlates with lower local polymorphism density (Col×Mh 27.1 polymorphisms/kb versus Col×Ler 80.7 polymorphisms/kb) in the 1,512 bp region spanning the peak intergenic marker (45 cM/Mb) until *GDSL*. Surprisingly, this region includes crossovers within the TE fragments (Col×Mh 8/124 and Col×Ler 5/240) (Fig. S3 and Table S13). We previously found that *RAC1* contains a strong CTT motif at its 5’-end, which have been associated with high crossover frequency (Choi *et al*, 2016, 2013; Shilo *et al*, 2015; Wijnker *et al*, 2013). Ler and Wl share a SNP in this motif but this does not obviously associate with differences in recombination rate [Col/Mh: CTTCGTCATCTTCTTCT; Ler/Wl: CTTCTTCATCTTCTTCT] (Fig. 3A).

### *RAC1* crossover frequency is resistant to changes in meiotic recombination pathways

Previous work has revealed an influence of interhomolog polymorphism on meiotic recombination pathways in Arabidopsis (Ziolkowski *et al*, 2015; Serra *et al*, 2018; Fernandes *et al*, 2017b; Girard *et al*, 2015; Emmanuel *et al*, 2006). Therefore, we sought to investigate *RAC1* crossover frequency in backgrounds with altered meiotic recombination pathways. Specifically we tested, (i) mutations in the anti-recombinases *recq4a recq4b*, *fancm* and *figl1* (Séguéla-Arnaud *et al*, 2015; Fernandes *et al*, 2017b; Crismani *et al*, 2012; Girard *et al*, 2015; Séguéla-Arnaud *et al*, 2016), (ii) mutations in the *msh2* MutS homolog (Emmanuel *et al*, 2006) and (iii) transgenic lines with additional *HEI10* copies (Ziolkowski *et al*, 2017). Each of these genotypes is available in Col and Ler backgrounds, which could be crossed together to generate Col/Ler F_1_ hybrids used for *RAC1* pollen-typing. We measured *RAC1* crossover frequency via DNA titration experiments (Fig. 5 and Tables S14-S17). The mean number of crossovers and parental molecules per μl were used to test for significant differences, by constructing 2×2 contingency tables and performing Chi-square tests. We compared three biological replicates of wild type Col/Ler F_1_ hybrids using this method, which did not show significant differences (Fig. 5A-5C and Tables S14-S17). Previous findings have demonstrated genome-wide crossover increases in hybrid *recq4a recq4b* and *figl1* mutants (Serra *et al*, 2018; Fernandes *et al*, 2017b; Séguéla-Arnaud *et al*, 2015), whereas *fancm* increases strongly in inbred, but not in hybrid backgrounds (Girard *et al*, 2015; Ziolkowski *et al*, 2015). Despite genome-wide crossover increases in these backgrounds, we observed that *RAC1* genetic distance significantly decreased in both the anti-recombination single mutants *recq4a recq4b*, *fancm*, *figl1* and *msh2*, and the multiple mutants *recq4a recq4b fancm* and *figl1 fancm* (Fig. 5A-5C and Tables S14-S17). Furthermore, when we compared wild type with lines containing additional *HEI10* copies we did not observe a significant difference in *RAC1* crossover frequency (Fig. 5D and Tables S14-S17). Therefore, in backgrounds with either increased class I (*HEI10*) and class II (*fancm*, *figl1*, *recq4a recq4b*) crossovers, the *RAC1* hotspot is unexpectedly resistant to increasing recombination frequency.

**Figure 5.**
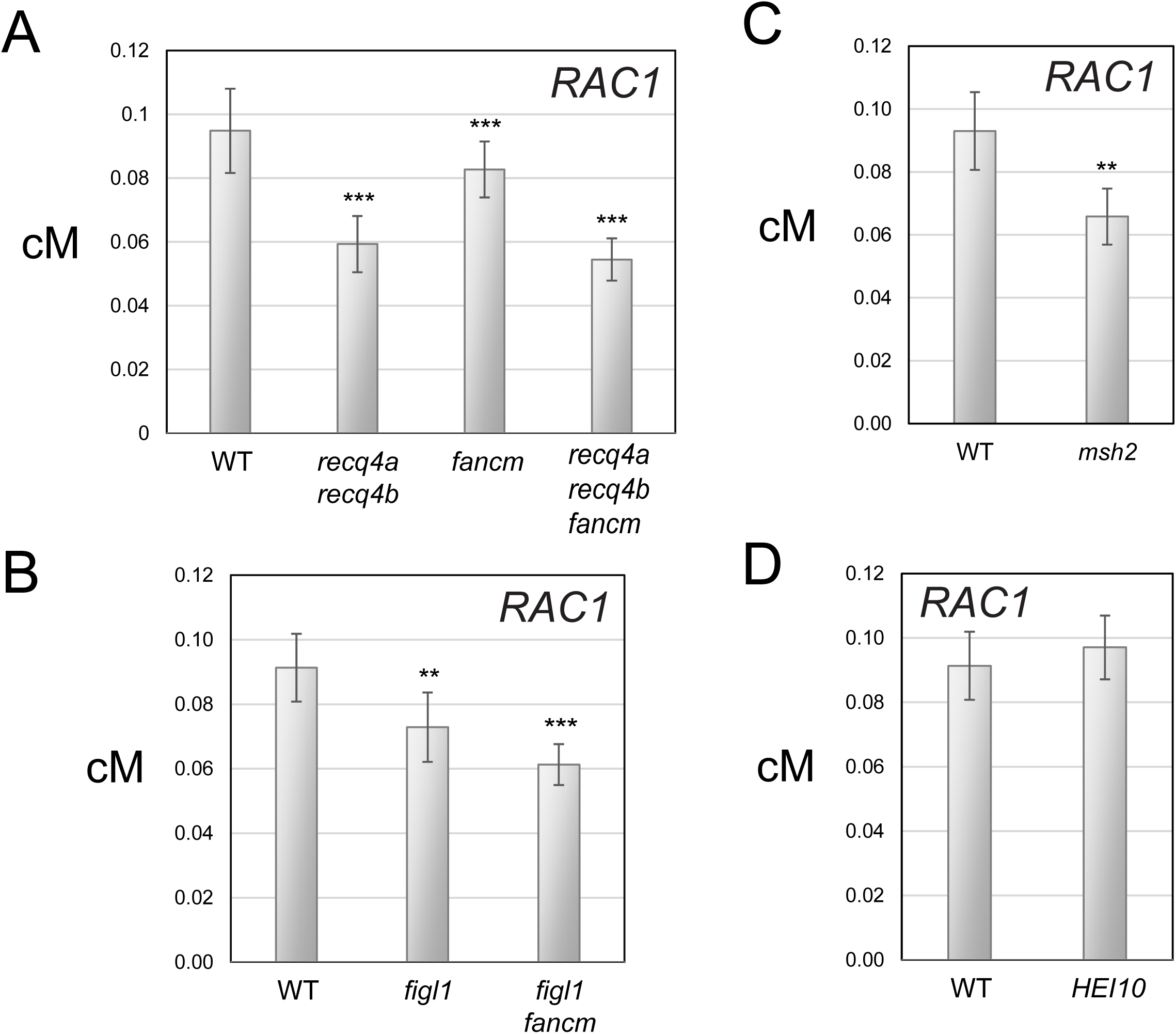
*RAC1* genetic distance in backgrounds with altered meiotic recombination pathways. **(A)** Barplots showing *RAC1* genetic distance (cM) measured in wild type, *recq4a recq4b*, *fancm* and *recq4a recq4b fancm* using single crossover and parental molecule titration. Error bars represent measurement standard deviation (square root of the variance). To test for differences the mean number of crossovers and parental molecules per μl were used to construct 2×2 contingency tables and Chi-square tests performed. The significance indicators ** and *** report a *P-*value of between 0.01-0.0001 and <0.0001, respectively. **(B)**. As for (A), but showing data for wild type, *figl1* and *figl1 fancm*. **(C)** As for (A), but showing data for wild type and *msh2*. **(D)** As for (A), but showing data for wild type and *HEI10*.

### *RAC1* crossover topology in *fancm* and *recq4a recq4b* anti-recombinase mutants

To analyse *RAC1* crossover distributions in wild type versus *fancm*, *recq4a recq4b* and *recq4a recq4b fancm* anti-recombinase mutants, we mass amplified crossovers and performed pollen-seq (Choi *et al*, 2017, 2016). In this approach, allele-specific PCR amplification is performed using multiple independent reactions seeded with an estimated ∼1-3 crossovers per reaction (Fig. 2A). Crossover concentrations are first estimated using previous titration experiments (Fig. 5 and Table S14). Mass amplified allele-specific PCR products are then pooled, sonicated and used for sequencing library construction (Choi *et al*, 2017, 2016). These libraries are subjected to paired-end 2×75 bp read sequencing (Table S18). We generated two biologically independent libraries for each genotype, sampling either ∼300 or ∼1,000 crossovers (Fig. 6A). The wild type 1,000 crossover dataset was previously reported (Choi *et al*, 2016).

**Figure 6.**
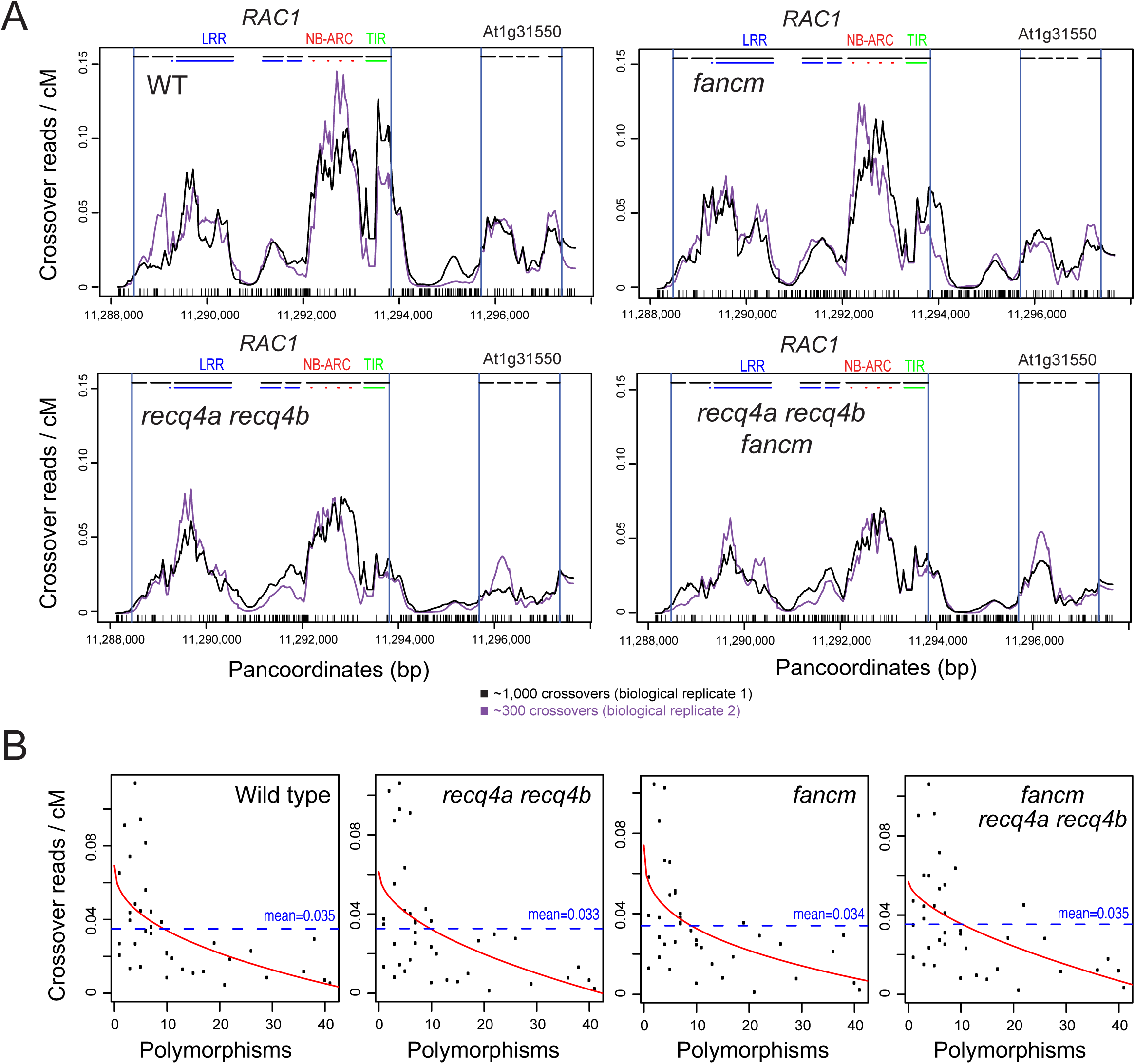
Crossover frequency across *RAC1* in wild type and *fancm*, *recq4a recq4b* and *fancm recq4a recq4b* anti-recombinase mutants. **(A)** The coverage of crossover reads aligned against the Col×Ler panmolecule was calculated and normalized by the total number of reads analysed. These data were also normalized by *RAC1* genetic distance (cM), measured previously by titration. For each genotype two biological replicates were analysed, constructed with amplifications from an estimated ∼300 (purple) or ∼1,000 (black) independent crossover molecules. Col×Ler polymorphisms are indicated by black ticks on the x-axis. Gene TSS and TTS are indicated by vertical blue lines, and exons by horizontal black lines. **(B)** 250 base pair adjacent windows were used to calculate the values of crossover reads/cM and polymorphisms (SNPs=one, indels=length) and plotted. The fitted line (red) was generated using the the non-linear model y = log(a) + b×x^(-c). y is reads/cM, x is polymorphisms, a is the intercept and b and c are scale parameters. The dotted horizontal line indicates the mean level of crossover reads/cM within the analysed region.

The Col and Ler *RAC1* haplotypes from our laboratory strains were Sanger sequenced, in order to generate templates for aligning sequence data to. Read pairs were split and aligned to Col and Ler haplotypes separately using Bowtie allowing no mismatches (-v 0), such that BAM alignment files were obtained for the reads aligned to either Col or Ler (Langmead *et al*, 2009). We then filtered for read pairs where one member mapped distally to Col and the other member mapped proximally to Ler, on opposite strands. This mapping configuration was expected due to the allele-specific primer orientation used during pollen-typing amplification. Consistent with these read pairs representing crossover molecules, their width distributions are similar to that of the sonicated PCR amplification products, prior to adapter ligation (Fig. S4). The crossover reads were then matched to the Col/Ler panmolecule, and counts were added to intervening sequences. These values were then normalized by dividing by the total number of crossover read pairs per library. Finally, the profiles were weighted by *RAC1* genetic distance (cM), measured previously via DNA titration (Table S14).

Overall recombination topology was similar between wild type and the anti-recombinase mutants *fancm*, *recq4a recq4b* and *recq4a recq4b fancm* (Spearman’s wild type vs *fancm r*=0.923 <2.2×10^-16^; wild type vs *recq4a recq4b r*=0.902 <2.2×10^-16^; *recq4a recq4b fancm r*=0.925 <2.2×10^-16^) (Fig. 6A, Fig. S5 and Table S15). Crossovers occurred mainly within the gene transcribed regions and were reduced in the highly polymorphic intergenic regions, in all genotypes (Fig. 6A and Fig. S5). In wild type, highest crossover frequency was observed at the *RAC1* 5’-end, with distinct peaks associated with the first and second exons, in addition to elevated crossovers occuring within the last three LRR domain-encoding exons (Fig. 6A and Fig. S5). Crossovers were also evident at the 5’ and 3’ ends of the adjacent gene (*GDSL*), although at a lower level to those observed in *RAC1* (Fig. 6A and Fig. S5). In *fancm* the crossover profile was similar, except for in the first *RAC1* exon where crossover frequency was reduced compared to wild type (Fig. 6A). In the *recq4a recq4b* and *fancm recq4a recq4b* mutants we observed that the *RAC1* 5’ crossover peaks in exons 1 and 2 were both reduced (Fig. 6A). The *RAC1* LRR crossover peaks in *recq4a recq4b* and *fancm recq4a recq4b* backgrounds were also less broad and became focused towards the end of exon 5 (Fig. 6A). The 5’-end of *GDSL* was also reduced in the *recq4a recq4b* and *fancm recq4a recq4b* mutants (Fig. 6A).

To investigate the relationship between crossovers and polymorphisms, we calculated recombination (crossover read pairs/cM) and polymorphism vlaues in 250 bp windows, against the *RAC1* Col/Ler panmolecule. Consistent with our previous observations, all genotypes showed a significant negative correlation between crossovers and polymorphism levels (250 bp adjacent windows, Spearman’s: WT *r*=-0.636, *fancm r*=-0.580, *recq4a recq4b r*=-0.567, *fancm recq4a reqc4b r*=-0.568) (Table S19). As described above, a non-linear model fitted the data using the formula y = log(a)+b×x^(-c), where y is the crossovers, x is polymorphisms, a is the intercept and b and c are scale parameters (Fig. 6B). Hence, the suppressive effect of polymorphisms was observed at *RAC1* in both wild type and anti-recombinase mutant backgrounds.

## DISCUSSION

The concentration of meiotic DSBs and crossovers into narrow hotspots is widespread among eukaryotes, which has important implications for genetic diversity and adaptation (Kauppi *et al*, 2004; Choi & Henderson, 2015; Baudat *et al*, 2013). For example, sequencing of SPO11-oligonucleotides has revealed meiotic DSB hotspots in fungi, animals and plants (Pan *et al*, 2011; Lange *et al*, 2016; Fowler *et al*, 2014; Choi *et al*, 2018). However, varying genetic and epigenetic factors control DSB hotspot location in these species. SPO11-oligo hotspots in budding yeast and plants are highest in nucleosome-free regions associated with genes and transposons (Pan *et al*, 2011; Sasaki *et al*, 2013; Choi *et al*, 2018; Wu & Lichten, 1994), which demonstrates the importance of chromatin for initiation of meiotic recombination. Variation in nucleosome occupancy and SPO11-1-oligos in plants also correlates with AT-sequence richness (Choi *et al*, 2018), which is known to exclude nucleosomes (Segal & Widom, 2009). In fission yeast SPO11-oligo hotspots are broader, located intergenically and do not show a clear association with nucleosome occupancy (Fowler *et al*, 2014). Mammalian meiotic DSB hotspots are directed to specific DNA sequences by binding of the PRDM9 KRAB-zinc finger protein (Baudat *et al*, 2013; Lange *et al*, 2016; Brick *et al*, 2012). PRDM9 possesses a histone methyltransferase SET domain which directs H3K4me3 and H3K36me3 histone modifications to nucleosomes flanking the bound DNA target sites (Baker *et al*, 2014; Lange *et al*, 2016; Yamada *et al*, 2017; Powers *et al*, 2016). Hence, depending on the species, chromatin and DNA sequence make varying contributions to the levels and fine-scale distributions of meiotic DSBs.

In many species, including budding yeast and plants, there is a positive correlation between meiotic DSB levels and the frequency of crossovers at the chromosome scale (Choi *et al*, 2018; Pan *et al*, 2011). However, extensive variation in the ratio of DSBs to crossovers is observed along chromosomes (Pan *et al*, 2011; Lange *et al*, 2016; Fowler *et al*, 2014; Choi *et al*, 2018). An extreme example occurs in fission yeast, where an inverse relationship is observed, with DSB hotspots occurring in regions of lowest crossover formation (Fowler *et al*, 2014; Hyppa & Smith, 2010). Variation in the crossover:non-crossover ratio has also been observed between mammalian hotspots (Kauppi *et al*, 2004; Cole *et al*, 2010; Baudat & de Massy, 2007; Jeffreys *et al*, 2001). A dramatic example of crossover:non-crossover ratio variation occurs at heterochiasmic mouse hotspots, where DSBs occur in both male and female meiosis, but crossovers only form in male meiosis (de Boer *et al*, 2015). Hence, the levels of initiating DBSs are important for crossover levels, but they are not the sole determinant of recombination outcome.

At *RAC1* and *RPP13* we observe a strong non-linear, negative relationship between interhomolog polymorphism and crossover formation. Hence, polymorphisms may contribute to variation in crossover:non-crossover ratios, by suppressing crossover maturation downstream of DSB formation. This is consistent with observations made in budding yeast where progressive addition of SNPs at the *URA3* hotspot reduced crossovers, with a simultaneous increase in noncrossovers (Borts & Haber, 1987). A likely mechanism for these effects is via MutS related heterodimers, including MSH2, which are capable of recognising mismatches and promoting dissolution of strand invasion events (Kolas *et al*, 2005; Alani *et al*, 1994; Hunter *et al*, 1996). Indeed, evidence exists in Arabidopsis for MSH2 acting as a hybrid-specific anti-recombinase at the megabase scale (Emmanuel *et al*, 2006). The inhibitory effect of interhomolog polymorphism on crossover formation may also account for discrepancies observed between SPO11-1-oligos and crossovers at the fine-scale (Choi *et al*, 2018). For example, at *RPP13* and *RAC1* we observe that SPO11-1-oligos are highest in nucleosome-depleted promoter, terminator and introns, whereas crossovers were highest in nucleosome occupied exons. It is possible that the mobility of strand invasion intermediates (Allers & Lichten, 2001), and repeated cycles of strand invasion during repair at single DSB sites (Oh *et al*, 2008; De Muyt *et al*, 2012), may also contribute to divergence between the location of initiating DSBs and the final crossovers. The phenomenon of crossover interference, which reduces the likelihood of adjacent DSBs being repaired as crossovers in the same meiosis, is also important to consider (Berchowitz & Copenhaver, 2010).

In addition to interhomolog polymorphism, chromatin marks may differentially influence meiotic recombination pathways and alter crossover:noncrossover ratios. For example, we observe that H3K4me3 is elevated at the 5’-ends of *RAC1*, *GDSL* and *RPP13*, which correlates with regions of high crossover activity. Although it is also notable that substantial 3’-crossovers also occur in these genes, where H3K4me3 occurs at lower levels. Although H3K4me3 levels do not strongly correlate with SPO11-oligo levels in animals, fungi or plants (Lange *et al*, 2016; Choi *et al*, 2018; Tischfield & Keeney, 2012; Pan *et al*, 2011), this mark is associated with recombination hotspots in multiple species (Shilo *et al*, 2015; Choi *et al*, 2013; Wijnker *et al*, 2013; Brick *et al*, 2012; Borde *et al*, 2009). In budding yeast the Spp1 subunit of the COMPASS methylase complex simultaneously interacts with H3K4me3 and the Mer2 meiotic chromosome axis component (Acquaviva *et al*, 2013; Sommermeyer *et al*, 2013), providing direct support for the tethered loop axis model (Blat *et al*, 2002). Analogous interactions are observed between mouse COMPASS CXXC1, PRDM9 and the IHO1 axis protein (Imai *et al*, 2017). Hence, the presence of H3K4me3 at the 5’-ends of *RPP13* and *RAC1* may promote crossover formation via similar mechanisms, downstream of DSB formation. Heterochromatic modifications also show specific interactions with the meiotic recombination pathways. For example, in Arabidopsis loss of CG context DNA methylation via the *met1* mutation, or loss of non-CG DNA methylation/H3K9me2 via *cmt3* or *kyp suvh5 suvh6*, both cause an increase in SPO11-1-oligos in pericentromeric heterochromatin (Choi *et al*, 2018; Underwood *et al*, 2018). However, the CG and non-CG mutants show increased and decreased pericentromeric crossovers, respectively (Choi *et al*, 2018; Underwood *et al*, 2018). This indicates that despite these heterochromatic mutants showing greater SPO11-1 DSB activity close to the centromeres, other chromatin features likely modify downstream repair choices.

In this study we measured *RAC1* crossover frequency in backgrounds with, (i) elevated *HEI10* dosage and thereby increased class I activity (Ziolkowski *et al*, 2017), (ii) increased class II crossovers via loss of function mutations in the *fancm, figl1* and *recq4a recq4b* anti-recombinases (Fernandes *et al*, 2017b; Crismani *et al*, 2012; Séguéla-Arnaud *et al*, 2016, 2015; Girard *et al*, 2015), and (iii) loss of function mutants in the mismatch repair factor *msh2* (Emmanuel *et al*, 2006). Despite these backgrounds showing elevations in crossover frequency, *RAC1* remained resistant to recombination increases or showed small but significant decreases. In this respect it is notable that genome-wide maps of crossovers in *HEI10*, *figl1*, *fancm* and *recq4a recq4b* backgrounds have shown that recombination increases are highly distalized towards the sub-telomeres, which are chromosome regions of lower interhomolog polymorphism (Serra *et al*, 2018; Fernandes *et al*, 2017b; Ziolkowski *et al*, 2017). Therefore, the location of *RAC1* on the edge of the chromosome 1 pericentromere may make this locus relatively insensitive to these distalized crossover changes. It is also possible that high polymorphism levels within *RAC1*, in addition to the surrounding regions of heterochromatin, may contribute to relatively stable crossover frequency between wild type and the high recombination backgrounds tested.

A local inhibitory relationship between polymorphism and crossover has implications for evolution of plant hotspots. Data from several species are consistent with meiotic recombination being mutagenic. This may occur as a result of DNA polymerase base misincorporation during DSB-repair accociated synthesis (Arbeithuber *et al*, 2015; Rattray *et al*, 2015; Yang *et al*, 2015), or mis-alignment during strand invasion causing insertions and deletions via unequal crossover (Sudupak *et al*, 1993). Therefore, high levels of recombination over many generations may cause higher levels of diversity and heterozygosity at hotspots, which may then suppress further recombination. Crossover inhibition is likely to be particularly potent when an unequal crossover generates large insertion-deletion polymorphisms, which are commonly observed at plant disease resistance loci (Choi *et al*, 2016; Sudupak *et al*, 1993; Kuang *et al*, 2008; Parniske *et al*, 1997). Despite the negative relationship that we observed between interhomolog polymorphism and crossovers at *RAC1* and *RPP13*, at the chromosome scale wild type crossovers in Arabidopsis show a weak positive relationship with interhomolog diversity (Serra *et al*, 2018; Fernandes *et al*, 2017b). Similarly, LD-based historical estimates are positively correlated with population diversity in Arabidopsis (Ziolkowski *et al*, 2015; Long *et al*, 2013; Cao *et al*, 2011). These population-scale relationships are likely to be partly explained by hitchhiking/background selection, causing more extensive reductions in diversity in regions of low recombination that are under selection (Barton & Charlesworth, 1998). Therefore, the relationship between interhomolog polymorphism and crossovers is complex, with both negative and positive relationships, depending on the species and scale analysed.

## MATERIAL AND METHODS

### Plant material

Arabidopsis lines used in this study were the *HEI10* line ‘C2’ (Ziolkowski *et al*, 2017), *recq4a-4* (Col, N419423) *recq4b-2* (Col, N511130) (Hartung *et al*, 2007), *recq4a* (Ler W387*) (Séguéla-Arnaud *et al*, 2015), *fancm-1* (Col, ‘roco1’) (Crismani *et al*, 2012), *fancm* (Ler, ml20), *figl1-1* (Col, ‘roco5’) (Girard *et al*, 2015), *figl1* (Ler, ml80) and *msh2-1* (Col, SALK_002708) (Leonard *et al*, 2003). Genotyping of Col *recq4a-4*, Col *recq4b-2*, Ler *recq4a* and *HEI10* T-DNA was performed as described (Serra *et al*, 2018). Col and Ler wild type or mutant backgrounds were crossed to obtain F_1_ hybrids, on which pollen-typing was performed. The *msh2-1* allele was introduced into Ler-0 background by six successive backcrosses. Genotyping of *msh2-1* was performed by PCR amplification using msh2-F (5’-AGCGCAATTTGGGCATGTCT-3’) and msh2-R (5’-CCTCCCATGTTAGGCCCTGTT-3’) oligonucleotides for the wild type allele, and msh2-F and msh2-T-DNA (5’-ATTTTGCCGATTTCGGAAC-3’) oligonucleotides for the *msh2-1* allele.

### *RPP13* and *RAC1* pollen-typing and Sanger sequencing

Pollen-typing was performed as described (Choi *et al*, 2017). Genomic DNA was extracted from hybrid F_1_ pollen (Col×Ler, Col×Wl or Col×Mh), and used for nested PCR amplifications using parental or crossover configurations of allele-specific oligonucleotide primers (Tables S20-S21). For each genotype replicate, ∼140 plants were grown and used for pollen collection. The relative concentrations of parental (non-recombinant) and crossover (recombinant) molecules were estimated by titration (Choi et al, 2017, Drouaud and Mézard, 2011; Kauppi et al, 2009). Recombination rate was calculated using the formula cM = (crossovers/(parentals+crossovers))×100. Amplified single crossover molecules were treated with exonuclease I (NEB, M0293) and shrimp alkaline phosphatase (Amersham, E70092), and then Sanger sequenced to identify recombination sites to the resolution of individual polymorphisms. For analysis we PCR amplified and sequenced the target regions from Col, Ler, Wl and Mh accessions, and used these data to generate Col×Ler, Col×Wl or Col×Mh panmolecules, which include all bases from both accessions (Fig. S2). To analyse the relationship between crossovers and polymorphisms we used adjacent 500 bp windows along the panmolecules and assigned crossover and polymorphism counts, where SNPs were counted as 1, and indels as their length in base pairs. When crossover events were detected in SNP intervals that overlapped window divisions the crossover number was divided by the proportional distance in each window. For example, if two crossovers were detected in a 150 bp interval, of which 50 bp were in window A and 100 bp in window B, we counted 2×(50/150) = 0.67 crossover in window A, and 2×(100/150) = 1.33 crossover in window B. A non-linear model was fitted to the data using the formula; y = log(a)+b×x^(-c), where y is the number of crossovers, x is polymorphisms, a is the intercept and b and c are scale parameters.

### *RAC1* mass crossover sequencing

Multiple independent *RAC1* crossover PCR amplifications were performed, where each reaction was estimated to contain between 1-3 crossover molecules, based on previous titration experiments. *RAC1* crossover amplification products were then pooled, concentrated by isopropanol precipitation and gel purified. 1-2 μg of DNA in 100 μl of TE was sonicated for each sample using a Bioruptor (Diagenode) (high setting, 30 seconds ON, 30 seconds OFF for 15 minutes), and fragments of 300-400 bp were gel purified, end-repaired and used to generate sequencing libraries (Tru-Seq, Illumina). The libraries were sequenced on an NextSeq instrument (Illumina) using paired-end 75 bp reads. Reads were aligned to the parental sequences (Col and Ler) using Bowtie, allowing only exact matches (Langmead *et al*, 2009). Reads were filtered for those that aligned to one parental sequence only. To identify crossover read pairs, we filtered for read pairs having a centromere-proximal match to Col and a centromere-distal match to Ler, on opposite strands, which is consistent with *RAC1* pollen-typing amplification. Read pair coordinates were then converted into pancoordinates using the Col/Ler key table (Table S2). A value of 1 was assigned to all panmolecule coordinates between each crossover read pair. This process is repeated for all read pairs and final values normalized by the total number of crossover read pairs, and finally weighted by genetic distance (cM).

## Data Access

The fastq files associated with mass crossover sequencing have been uploaded to ArrayExpress accession E-MTAB-6333 (“Meiotic crossover landscape within the *RAC1* disease resistance gene”).

## Acknowledgements

We thank Raphael Mercier for provision of *fancm*, *figl1* and *recq4a recq4b* mutants in the Ler background (IJPB, INRA, France). Research was supported by a Royal Society University Research Fellowship, the Gatsby Charitable Foundation grant GAT2962, BBSRC grant BB/N007557/1, BBSRC grant BB/L006847/1, BBSRC-Meiogenix IPA grant BB/N007557/1, ERC ‘SynthHotSpot’ Consolidator Grant, National Natural Science Foundation of China grant 61403318, RDA Next-Generation BioGreen 21 Program PJ01337001, NRF Basic Science Research Program NRF-2017R1D1AB03028374, EMBO long-term postdoctoral fellowship ALT 807-2009, the Bettencourt Schueller Foundation and a Gatsby Ph.D studentship.

